# Preparation of large biological samples for high-resolution, hierarchical, multi-modal imaging

**DOI:** 10.1101/2022.07.02.498430

**Authors:** J. Brunet, C. L. Walsh, W. L. Wagner, A. Bellier, C. Werlein, S. Marussi, D. D. Jonigk, S. E. Verleden, M. Ackermann, Peter D. Lee, Paul Tafforeau

## Abstract

Imaging the different scales of biological tissue is essential for understanding healthy organ behavior and pathophysiological changes. X-ray micro-tomography using both laboratory (µCT) and synchrotron sources (sCT) is a promising tool to image the 3D morphology at the macro- and micro-scale of large samples, including intact human organs. Preparation of large samples for high resolution imaging techniques remains a challenge due to limitations with current methods, such as sample shrinkage, insufficient contrast, movement of the sample and bubble formation during mounting or scanning. Here, we describe a protocol to prepare, stabilize, and image large soft-tissue samples with X-ray microtomography. We demonstrate the protocol using intact human organs and Hierarchical Phase-Contrast Tomography (HiP-CT) imaging at the European Synchrotron Radiation Facility, but the protocol is equally applicable to a range of biological samples, including complete organisms, for both laboratory and synchrotron source tomography. Our protocol enhances the contrast of the sample, while preventing sample motion during the scan, even in case of different sample orientations. Bubbles trapped during mounting and those formed during scanning (in case of synchrotron X-ray imaging) are mitigated by multiple degassing steps. The sample preparation is also compatible with magnetic resonance imaging (MRI), CT, and histological observation. We describe a protocol for sample preparation and mounting which requires 25 to 34 days for a large organ such as an intact human brain or heart. This preparation time varies depending on the composition, size, and fragility of the tissue. Use of the protocol enables scanning of intact organs with a diameter of 150 mm with a local pixel size of one micron using HiP-CT.

## Introduction

Quantifying the morphology of our organs in health and disease is a complex imaging task involving multiple scales and often modalities. Characterising the interactions between each of these scales is essential. However, most imaging techniques are limited by either resolution or field of view, and bridging the macro- to micro-scale remains a challenge. Classical methods such as visualisation through serial sections with conventional histology^1–3^ or electron microscopy^4–6^ provide quantification of the tissue’s microstructural organisation and composition; however, these techniques normally require sectioning of the organ into smaller samples, and are extremely labour intensive and time consuming. Although very promising, the same drawbacks apply to optical clearing combined with light-sheet microscopy^7,8^. Methods like optical coherence tomography^9,10^, multiphoton microscopy^11^, or confocal microscopy^12,13^ can capture the local 3D microstructure of the tissue at the cellular scale, but their limited tissue penetration hinders the deep tissue imaging required to observe entire, intact organs^14^. Recently, high-resolution MRI achieved a voxel size of 100 µm in an whole ex vivo human brain^15^. Although this imaging technique is non-destructive and has a large field of view^16^, this resolution is still not sufficient to examine the microstructure. Hierarchical imaging are techniques capable of overcoming the trade-off between resolution and field of view. In this approach, multiple images of the same sample are acquired at different resolutions to bridge the different scales. In Verleden et al.^17^, laboratory µCT has been used to image entire lungs with 150 micron voxels, followed by subsequent local sample cores extraction; these cores were then scanned with µCT to achieve 10 micron voxels.

Considerable progress has been made in the field of X-ray imaging over the last decade, especially in soft tissue visualisation^18^. Synchrotron X-ray tomography (sCT) has proved to be one of the most powerful X-ray-based imaging techniques due to its high brightness^19^, enabling the observation of soft tissue at high resolution, with enough contrast to see the structural components^20–22^. In particular, phase-contrast-based sCT^23^, combined with the right sample preparation and mounting procedure has provided extensive information on biological tissues such as heart fibre orientation^24,25^ or brain cellular map^26^. Phase-contrast enables visualisation of soft tissues that would be invisible with conventional X-ray tomography without the use of a contrast agent^27^. Nevertheless, the studies using sCT are mostly limited to organs of fetus^28^, small subsamples of human organs^29^, or organs from small animal models, e.g. mice^30^ and rabbit^31^, due to the restricted field of view. Some studies have imaged larger samples, for instance Dutel et al.^32,33^ observed coelacanths with a diameter reaching 10 cm and a height of 30 cm; however, the voxel size was limited to 30 µm and the structure of interest were hard tissues.

New 4^th^ generation synchrotron sources, such as the European Synchrotron Radiation Facility (ESRF)’s Extremely Brilliant Source (EBS), provide enough beam coherence and flux to visualise intact human organs from the macro- to the micro-scale. Using the ESRF-EBS we have recently developed a new technique, Hierarchical Phase-Contrast Tomography^34^ (HiP-CT) that allows scanning of large intact human organs with 25 µm voxels, with subsequent zooming (without sectioning), achieving one micron voxels locally. Although often overlooked, sample preparation and mounting are crucial to achieve the highest resolutions with this and other techniques, especially when visualising large soft tissue samples, such as human organs, where the ratio of voxel size to organ diameter is 1:150,000 (1 µm in 150 mm).

The success of all these experimental techniques depends on careful preparation of the sample to avoid artefacts in the images. Soft tissue imaging using µCT or sCT presents a number of challenges compared to hard tissues. One of them is the lack of contrast when imaged with X-rays due to the similar densities of the sample components. Furthermore, the back-projection algorithms reconstructing the 3D volume from X-ray projections assume no movement of the sample during scanning. Most sample preparation methods are designed to prevent drifting or deformation of the sample to avoid movement artefacts that would reduce the quality of the images^35^. Some reconstruction algorithms have been developed to overcome this issue^36^ but they remain complex and computationally costly to operate.

Although the protocol we describe is applicable to a wide range of biological sample over a range of sizes and imaging modalities, this paper will focus on the example of synchrotron phase contrast imaging. The benefits of the different steps will vary depending on both sample size and imaging modality.

### Development and overview of the protocol

The sample preparation and mounting protocol described herein was developed from the need to image large biological volumes, such as intact human organs. In our recent publication^34^, we presented a new imaging technique called Hierarchical Phase-Contrast Tomography (HiP-CT) using the ESRF’s EBS to study five intact human organs: lung, brain, heart, kidney, and spleen. This technique enables scanning of intact organs with 25µm voxel size. Areas are then selected for further high-resolution scanning without the requirement for biopsies. To image such large structures, a high-energy X-ray beam is required to penetrate the samples. We used a polychromatic beam with energy ranging from 64 to 80 keV for organ imaging, but higher energies (up to 140 keV) would be more adapted if available.

Preparation and stabilisation of the organ for this technique is essential as any density inhomogeneity, remaining gas bubbles, or movement during scanning would greatly reduce the image quality. During the development of the method, several problems arose. The first issue was related to bubble trapping during mounting, and bubble formation during the scan, creating both motion blurring and phase contrast artefacts, reducing scan quality. This was resolved by limiting the dose absorbed by the sample and including multiple degassing steps during the mounting procedure. As the sample is large, the scans require a long time to complete, thus maintaining the specimen in a fixed position is essential. This challenge was solved by using a mixture of crushed agar gel and liquid (in our case ethanol 70%) as a mounting media around the organ. In addition, the dehydration of the organ with ethanol increased the contrast of the images^37^ and diminished the bubble formation.

Here, we present a procedure to prepare whole human organs for imaging with sCT, µCT, medical CT and MRI, that is compatible with a final stage of classical paraffin embedded histology. In this protocol, we mainly describe the sample preparation and mounting with ethanol-agar; the X-ray imaging protocol using HiP-CT is detailed in our recent methods paper^34^, while its application to a medical challenge (quantifying the damage COVID-19 does to lung vasculature) is explored in an applications paper^38^. In brief, after fixation of the body (Steps 1A(i)), the organ is extracted (Step 1A(ii)-1A(iii)), immersion fixed (Step 2), dehydrated and degassed with vacuum (Step 4A) or thermal cycles (Step 4B) depending on its fragility, mounted with crushed agar gel mixture (Steps 5-32), and imaged using sCT, µCT, clinical CT, and MRI (Step 33), finally, an histological analysis is performed (Step 34). See Figure 1 for an overview of the procedure.

**Figure 1:**
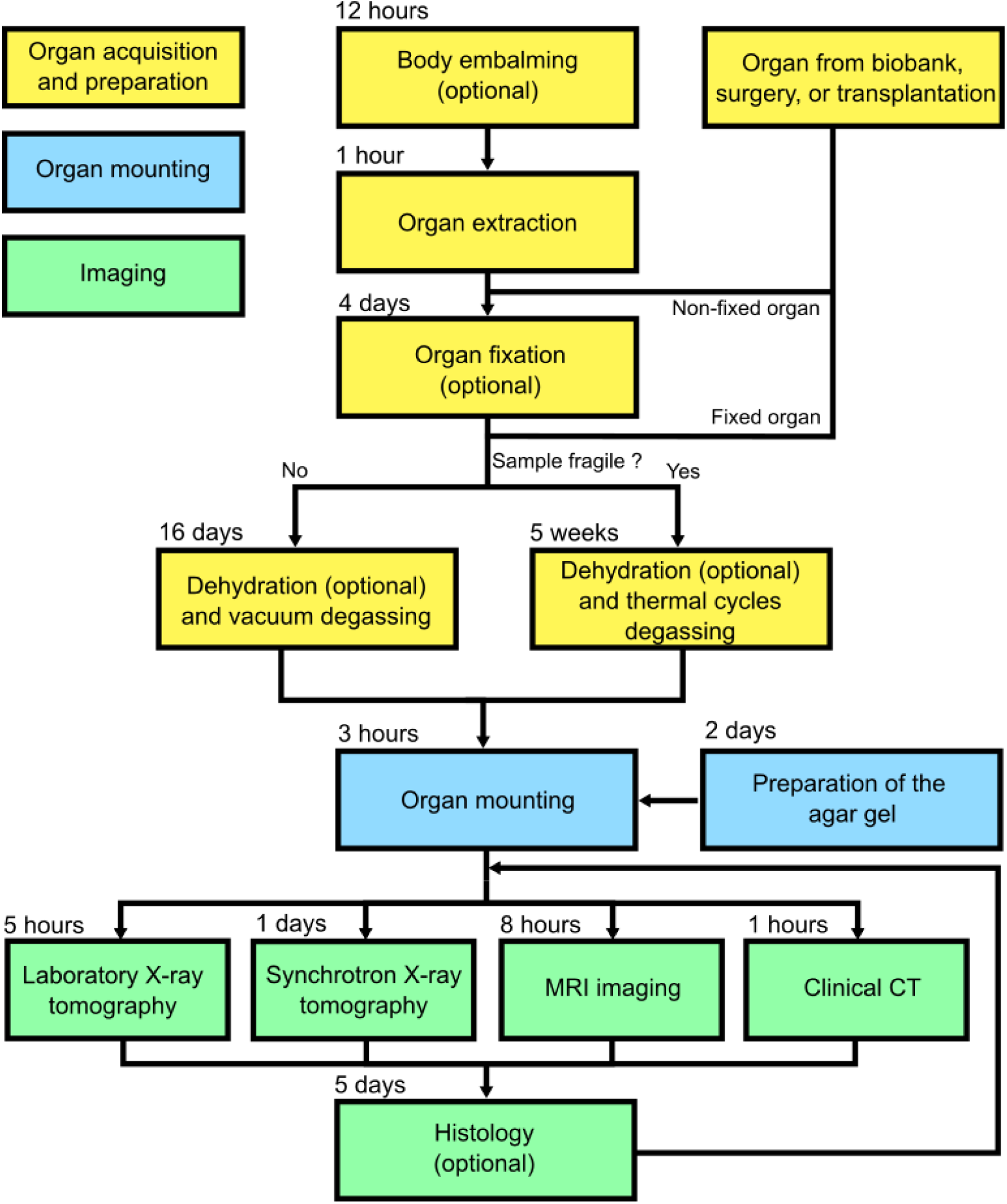
Overview of the sample preparation, stabilization, and scanning of large biological sample. The protocol contains three major steps (organ acquisition and preparation, organ mounting, and imaging) indicated in color-coded boxes. The organ can be retrieved either from a biobank, a surgery, a transplantation, or it can be extracted from a donated body. We provide two protocols for degassing depending on the fragility of the organ (vacuum degassing and thermal cycles degassing). Once the sample is mounted with the agar gel, different imaging techniques can be performed (µCT, sCT, clinical CT, and MRI). After imaging, histology can be carried out on the sample. The times for each steps are based on Walsh et al.^34^.

### Advantages of the protocol compared to other methods

This protocol was developed and optimized to image intact human organs at high-resolution with HiP-CT, as shown in FigureFigure **2**, and has been applied to various organ types including brain, heart, lung, kidney, and spleen. Nevertheless, the organ preparation procedure is flexible and can be used with animal organs, soft tissue in general, or even on complete small animals.

**Figure 2:**
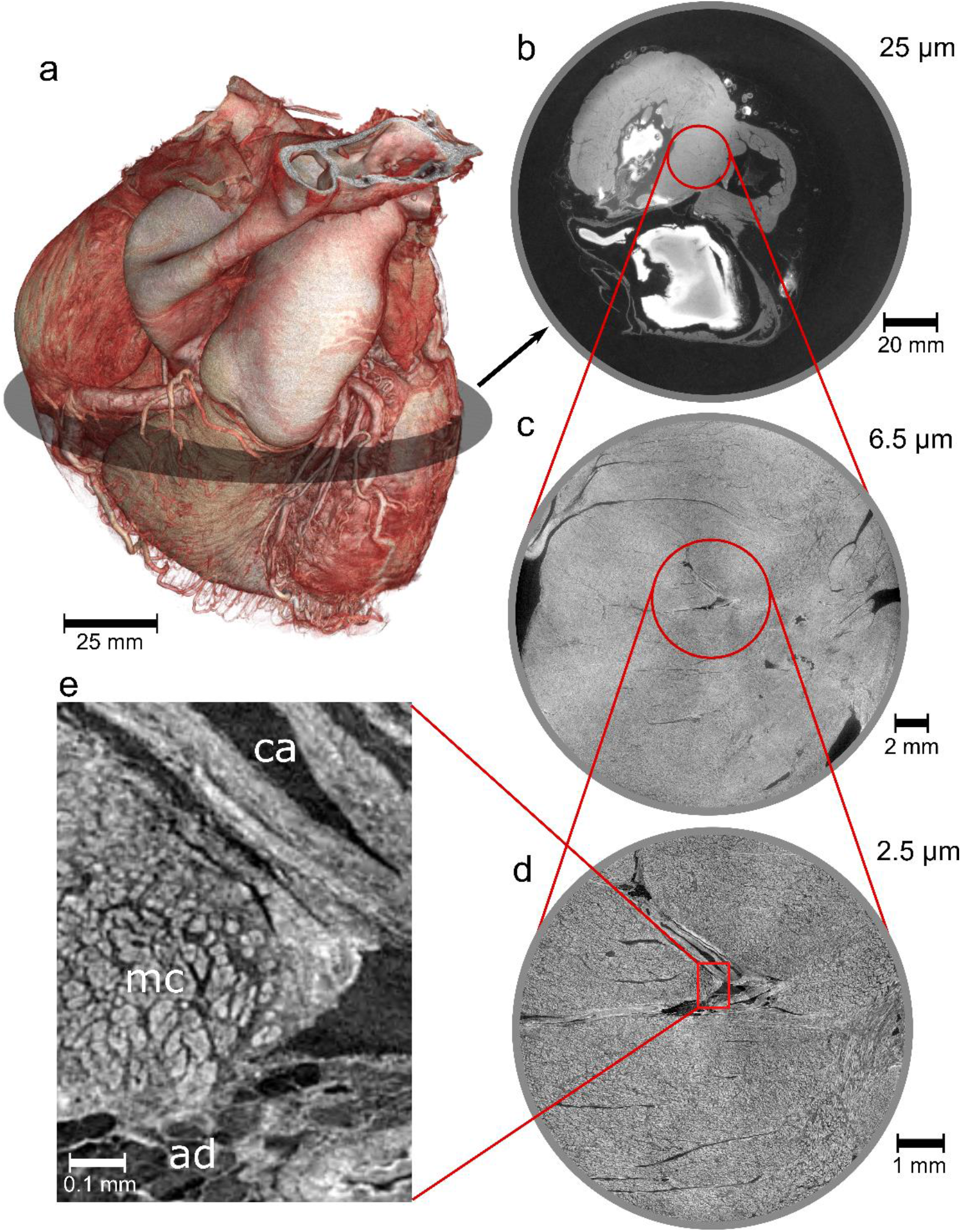
HiP-CT images of an intact human heart. **a**, 3D view of the human heart. Cross-sections of the heart with a voxel size of 25 µm (**a**), 6.5 µm (**c**), and 2.5 µm (**d**). **e**, Magnification of the 2.5 voxel image with annotation on the principal structures observed (ca, coronary artery; mc, myocyte cells; ad, adipose tissue). All experiments followed the relevant governmental and institutional ethics regulations for human experiments.

Most common sample preparation methods for X-ray imaging include fixation, alcoholic dehydration, paraffin-embedding, or critical-point drying. One of the main drawback of wet embedding such as alcoholic-immersion is sample drif which creates movement artefacts^35^. In our protocol, this is prevented by the use of small blocks of agar gel in density equilibrium with the mounting liquid, holding the sample in place. In Walsh et al.^34^, it is shown that scans taken several months apart on the same sample prepared with our organ-stabilization protocol can simply be registered using manual rigid transformation. This demonstrates the stability of this method over long time periods. An issue common to all these sample preparation methods is tissue shrinkage. This effect can hinder morphological quantification and lead to erroneous analyses. However, compared to paraffin embedding^39,40^ and critical point drying^35,41^, the sample shrinkage can be mitigate using multiple ethanol baths in ascending ethanol concentrations^37^ up to 70%, thus preserving the morphology of the tissue. Furthermore, the dehydration of the sample with ethanol ensures a higher contrast with µCT^37,42^ or sCT^43^. This specimen preparation protocol is compatible with MRI, clinical CT, and histology (after 3D imaging and dismounting). The protocol can be applied to large and small biological samples and multiple synchrotron beamlines could benefit from it, such as I13-2 at diamond, Tomcat at Swiss Light Source (SLS), or 20XU at Spring 8. Finally, this technique is easy to implement as the materials are readily available and it requires simple equipment.

### Limitations

Although this procedure presents several advantages compared to existing sample preparation protocols, some limitations should be noted.

#### Fixation and dehydration of the sample

The timing of the different steps of the protocol involving fixation and degassing are heavily dependent on the composition and size of the tissue. In the present procedure, we give examples of timing for different human organs; however, timing for other type of tissues may vary. Furthermore, multiple steps of the protocol could be further optimised.

Whilst fixation of the organ is critical for long term tissue preservation, it heavily affects tissues mechanical^44,45^, diffusion properties which could confound other measurements. Despite increasing the image contrast, ethanol dehydration also affects the mechanical properties of the tissue^37,46^. This limits the use of this preparation method for in situ testing; however, by not fixing the tissue and replacing ethanol with water, the protocol could be used to quantify mechanical properties over a short duration (as biological degradation would occur). If the contrast provided by ethanol is not high enough, or not adapted to the needs, various contrast agent could be used. In some cases, other techniques like DiceCT may be more suitable^47^. If agar gel was not desirable for a particular application, it could be replaced with other solid or elastic media that would be in equilibrium with the mounting liquid and relatively amorphous in its structure. For instance, in case of mounting with 96% ethanol, transparent candle crystal gel can be used instead of agar gel. In case of mounting with water, gelatin blocks or polyacrylamide blocks can also be used.

#### X-ray dose limit

One of the main drawbacks of wet embedding methods is the formation of bubbles due to dose rate or dose accumulation in case of sCT^48^. These bubbles can move or damage the sample, in addition they can create strong artefacts dramatically reducing the image quality^49^. Although bubbling is an issue with other standard preparation methods e.g. paraffin-embedding, it is not present with critical-point drying. In our protocol, this problem is mitigated by multiple degassing steps during the procedure, delaying the nucleation of bubbles during the scan. This problem is also mitigated in our case by the use of high-energy X-rays, however only a few synchrotrons are equipped to image soft tissue at high energy, as this reduces contrast. For low-energy X-ray tomography, the degassing steps must be performed conscientiously, and the dose rate controlled.

### Experimental design

Here, we describe a protocol to prepare, mount, and scan large human intact organs, but the approach is not limited to human samples and could be expanded to large biological samples, such as large muscles, complete limbs or complete animals. The timing of the different steps detailed here has been optimized for human organs; however, depending on the composition and size of the organ the timing and quantity should be adapted. Experiments involving humans or animals must be carried out in accordance with institutional guidelines and laws, following protocols approved by local ethics committees.

#### Sample collection and fixation

Biological tissue can be collected by various methods. If the sample is collected directly from a biobank, surgery or transplantation (step 1B), it can be taken directly to the sample fixation stage (step 2). If the organs come from a donated body, organ extraction must be performed (step 1A(ii)). Embalming of the body is carried out shortly after death, prior to organ extraction (by a licenced practitioner). Here, we describe the method used in Walsh et al.^34^. The body is fixed by injecting formalin diluted in a solution containing lanolin into the right carotid artery (Step 1A(i)). After evisceration, complete fixation of the organ is ensured by immersing it in 4% neutral buffered formaldehyde (Step 2). The duration of fixation is defined by the size of the organ, e.g. 4 days for a human brain. When eviscerating the organ, ensure as much surrounding tissue as possible is removed to decrease the time of penetration of the fixative into the organ. The volume of fixative is also important, we recommend a volume at least 4 times greater than the tissue volume.

Some organs, such as the lung, may require inflation. This can be partially accomplished at the fixation stage by using instillation of formalin in the lungs under controlled pressure. Once fixated, the replacement of formalin by the successive baths of ethanol with the vacuum pumping is coupled with injection of ethanol in the bronchia to keep consistent 3D shape. Strictly controlled pressure with ethanol degassing is not be possible as the only way to achieve it is via using degassed ethanol instilled at a controlled pressure followed by clamping of the bronchia. This would lead to explosion of the lung if it was degassed while clamped. However the results Walsh et al.^34^ show that without strictly controlled inflation at the ethanol dehydration stage, the result is still has high scientific utility.

#### Organ dehydration and degassing

Here, the organ is dehydrated with multiple pre-degassed ethanol baths. The transition to 70% ethanol must be smooth enough to avoid shrinkage^37^. 70% ethanol concentration was chosen as the best compromise between controlled shrinkage and sufficient contrast to observe structures of interest using phase-contrast; however, a different final ethanol concentration can be used depending on the application. During this step, degassing has to be performed to remove free and dissolved gas present in the tissue in order to avoid bubble formation during imaging. For human organs, we provide two different methods depending on the fragility of the organ for dehydration and degassing. The vacuum degassing (Steps 4Ai-v) can be used for most organs (heart, lung, liver, kidney, spleen). In this method, the organ is immersed in 4 successive pre-degassed ethanol baths of concentration 50%, 60%, 70%, and then a second 70% bath to ensure equilibrium is reached. A degassing step is performed between each bath. Equilibrium is generally reached in 4 days for each concentration for a human organ such as a brain or a heart. Degassing is performed with a vacuum pump and a desiccator. Alternatively, thermal cycling (Steps 4Bi) was developed for fragile organs, such as human brain, as some damage was observed after using the vacuum degassing method^34^. In this method, 4 thermal cycles are performed by immersing the organ in four successive baths of 50%, 60%, 70%, and second 70% pre-degassed ethanol. Each thermal cycle consists of immersing the organ in the highly degassed ethanol bath at room temperature. The container has to be closed with special care not to entrap any bubbles. It is then kept in a fridge for 4 to 5 days at 4°C. During this period, the dissolved gas will diffuse in the surrounding ethanol, and the bubbles will progressively dissolve too. After this time, the solutions and organ must be brought back to room temperature, and a new cycle can be started using a new strongly pre-degassed ethanol bath. For both methods, the minimum number of ethanol baths is 4 to reach 70% of final concentration without having substantial shrinkage. The immersion times have to be adapted and optimized to the type and fragility of the organ. The result of the degassing can be tested by making a radiograph of the organ in its jar without the mounting media described hereafter. If some remaining bubbles are still visible, more thermal cycles can be performed with ethanol at 70%. Organs with adipose tissues, like the brain, require a longer time to equilibrate with ethanol. At each stage dehydration can be checked by disturbing the container with the organ inside and looking for streaks of different density (different transparency) forming in the surrounding ethanol solution, this indicates the presence of water in the ethanol.

Ethanol dehydration is sufficient to observe the structures of interest with µCT and sCT when using phase-contrast; however, it should be possible to combine this protocol with the use of a contrast agent to resolve specific components of the sample or when using less sensitive imaging techniques. The contrast agent must be miscible with 70% ethanol. Contrast agents would increase the absorption of the sample, which may enhance bubble formation during imaging if using intense synchrotron X-ray beam.

In cases where the characterization technique is not be compatible with ethanol (as for instance MRI imaging for diffusion), the same protocol for degassing and then for mounting can be applied using water with formalin at the desired concentration. Specific safety aspects (such as working under an adapted fumehood) have to be taken as vacuum degassing solutions of formalin can produce dangerous vapors.

#### Bubble formation

One of the main problems in soft tissue imaging in wet embedding is the presence of bubbles. These can come either from the mounting protocol (bubbles in the organs or trapped in the mounting media), from very high X-ray doses resulting in evaporation of the ethanol (in case of sCT). In both cases they can create significant artefacts (Figure 3), and in case of bubbling event due to the dose, the movements of the bubbles during the scanning often renders the scans unusable^50,51^. Preliminary tests showed that degassing the sample prior to imaging removes trapped bubbles and delays the apparition of new bubbles, most probably by delaying the nucleation effect of the bubbles. Hence, several degassing steps were incorporated in the protocol to mitigate the bubble formation.

**Figure 3:**
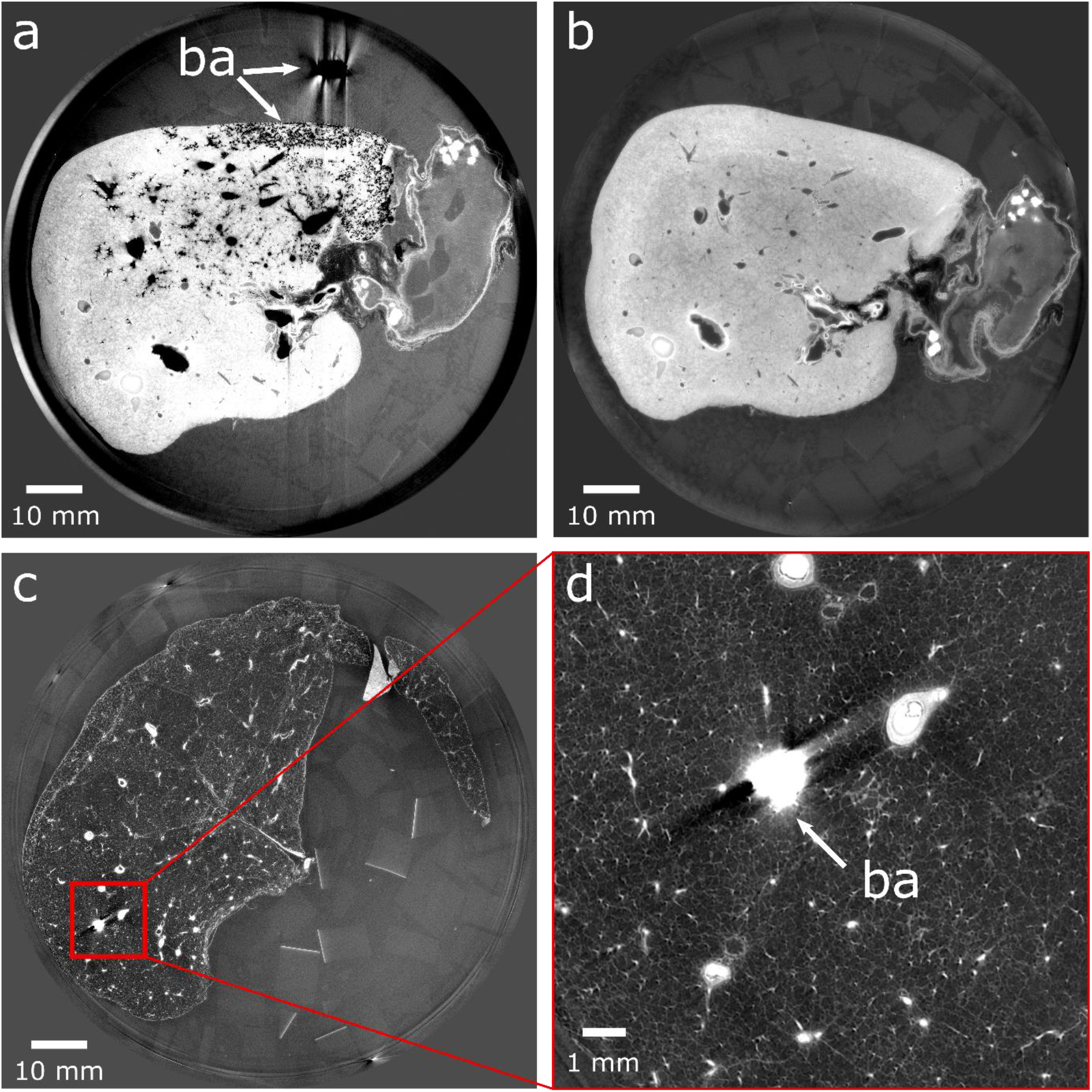
Example of bubble artefacts due to the radiation dose (a,b) or to a stable bubble entrapped during mounting (c,d). **a**, Cross-section of a human liver with numerous bubbles that have developed due to too high a dose uptake during scanning causing artefacts in the images (ba, bubble artefact). **b**, Cross section of the liver imaged after removal of bubbles by degassing the sample with vacuum degassing. **c**, Cross-section of a human lung with a bubble entrapped during mounting. **d**, Zoom on the bubble trapped in the lung vascular system. All experiments followed the relevant governmental and institutional ethics regulations for human experiments.

#### Organ mounting

The purpose of this step is to maintain the organ in position during the scan to avoid motion artefacts and to ensure that more scans can be performed at a later stage with good 3D rigid registration between subsequent experiments. The agar gel cannot be prepared with 70% ethanol directly, it has to be done by mixing agar-agar (20g/L) with demineralized water. Once gelled, the mixture can be cut into small cubes (step 8) and poured into 96% ethanol (6L:2L ethanol:agar-agar) (step 9), ensuring a final ethanol concentration of 70%. After degassing (step 12) and crushing of a part of the agar cubes (step 14), the mixture is ready for mounting. If a mounting with formalin is preferred to ethanol (e.g. to improve contrast with MRI), the 96% ethanol bath can be replaced with a 4% formalin bath instead. The agar gel can be prepared in advance and stored in an airtight container to prevent the introduction of gas in the solution. The amount prepared depends on the size of the organ and container used for final mounting. An agar gel prepared with 70% ethanol ensures a drastic diminution of bubble formation during imaging due to the reduced density of the solution. It is important to use a crushed agar gel and not a blended agar gel, as the blended gel does not hold the sample as firmly as the crushed agar gel and could lead to movement during imaging. Initially, only agar cubes were used to hold the sample in position; however, the cubes were found to be too rigid, creating deformations where they contacted the surface of soft organs such as lungs. Thus, the cubes should only be used at the bottom of the container, a few centimeters, to create a solid base and avoid rotation of the sample (step 16). Crushed agar is used in the remaining of the container to maintain the sample in position. The mix of crushed agar in the mounting liquid (70% ethanol or 4% formalin here) has to be added to the container gently with a ladle to avoid gas bubble entrapping during the process (Figure 4c). The dimensions of the container should be as close as possible to the specimen to minimise the amount of material that the X-ray beam has to pass through, however, the organ should not touch the side of the container to avoid artefacts in the images that would compromise the accuracy of the reconstruction. In case of sCT, the container used for the mounting must be made of a material resistant to X-rays, like Polyethylene terephthalate (PET). Once the agar gel has been compacted around the sample (step 22), the container can be sealed with a liquid-tight lid (step 28). The mounting should be assessed to ensure that no movement of the sample in the container is possible and no bubbles can be seen inside. If some bubbles are entrapped, rapid vacuum degassing can be used to remove them. The same approach can be used to remove bubbles in an organ in case of a bubbling event due to high dose during sCT scanning, (NB, do not degas the sealed container, remove the lid first then degas). An example of insufficiently compacted agar is shown in the online supplementary video 1. With the organ fixed and placed in 70% ethanol, the sample can be stored for years (Figure 4d).

**Figure 4:**
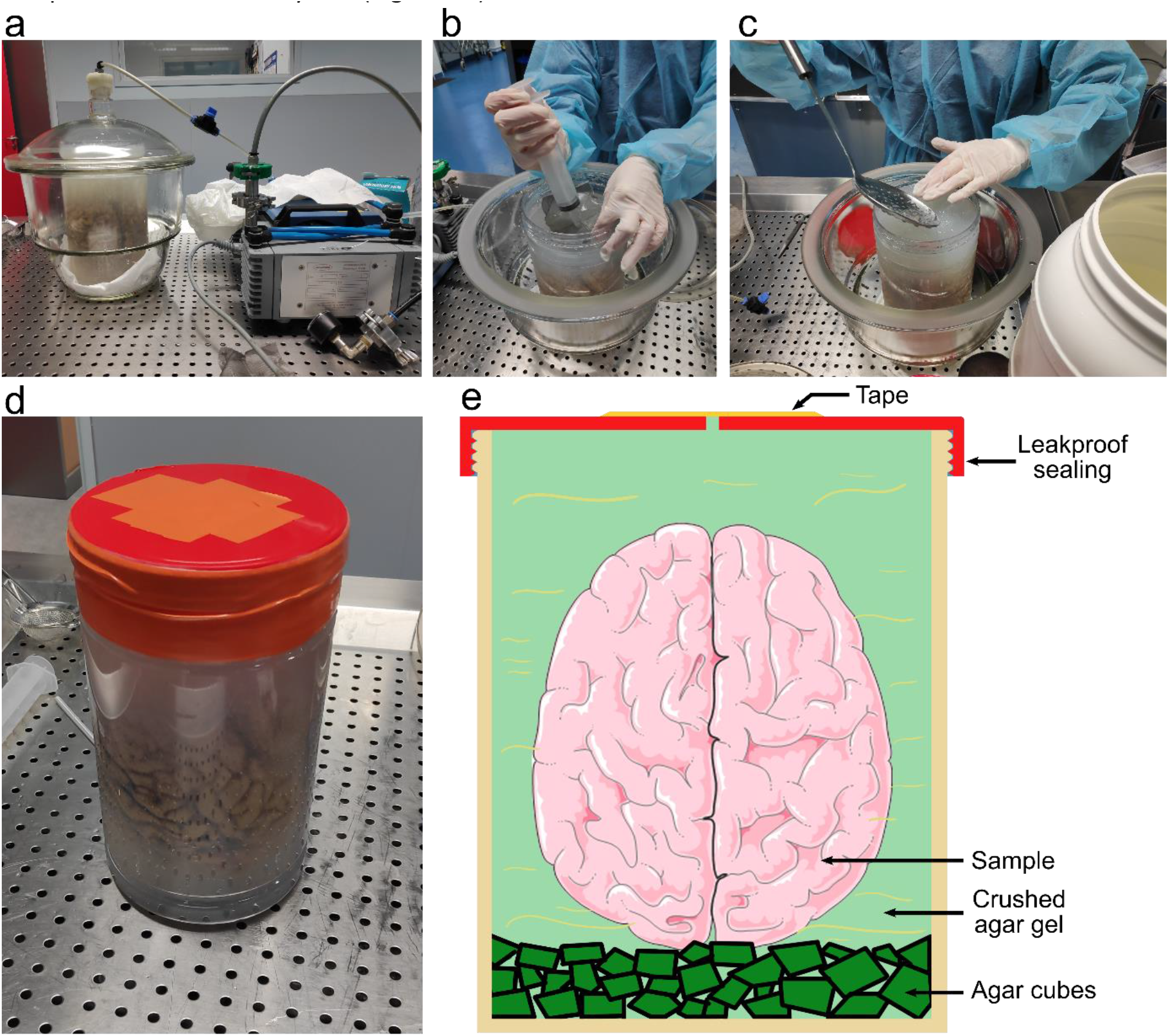
Procedures for organ mounting. **a**, Degassing of the organ in its container using a desiccator and a vacuum pump. **b, c**, After compression of the agar gel present in the container, some of the ethanol is removed using a syringe and a sieve (b), then agar gel is added (c). This process is continued until the agar gel is sufficiently compressed around the sample to hold it in position. **d**, Mounted sample stabilized and degassed in the agar gel, ready to be scanned. **e**, Schematics of a mounted brain, like in **d**, fixed with the crushed agar gel and agar cubes, in a sealed container. All experiments followed the relevant governmental and institutional ethics regulations for human experiments.

#### sCT imaging and reconstruction

The X-ray imaging and 3D reconstruction are not described in detail in this protocol; however, a detailed description is available in Walsh and al.^34^ or by contacting corresponding authors. This method was developed for large organ imaging with HiP-CT but many aspects would be beneficial for more classical sCT and uCT (laboratory source) imaging on smaller samples, including the immobilisation and contrast enhancement linked to ethanol preparation. Once the sample is mounted, scanning can be performed at any time, with typical sCT imaging results of an intact human heart shown in FigureFigure **2**. The main limitation of this technique, is the apparition of bubbles following high dose X-ray application to the organ in the case of sCT. If only few bubbles appear during imaging, the sample can be left in a refrigerator at 4°C to dissolve them again. If this is not successful, the sample can be re-degassed with vacuum pumping without having to dismount the sample, but there are inevitably some small movements during the process. In severe cases with too many bubbles to be removed by rapid vacuum degassing, or if scanning fragile samples such as brain, the mounting must be redone. Several solutions exist to avoid the apparitions of bubbles during sCT and concern the optimization of detection, the increase of relative contrast to work with lower dose, a better control of the dose, better data processing, or having resting period for the samples between successive scans of the same area. Bubbling seems to be strongly linked to the thermal effect of dose on the evaporation rate of the ethanol. Developing sample environmental chambers to work at lower temperature and ensuring a better thermal equilibrium may be an efficient way to alleviate the risk of bubbling linked to the use of 70% ethanol

#### MRI imaging

The sample preparation method presented in this protocol is compatible with MRI (Figure 5c), making it highly amenable to multi-modal studies. However, a number of considerations should be taken into account before deciding on the sample preparation method and scanning order.

**Figure 5:**
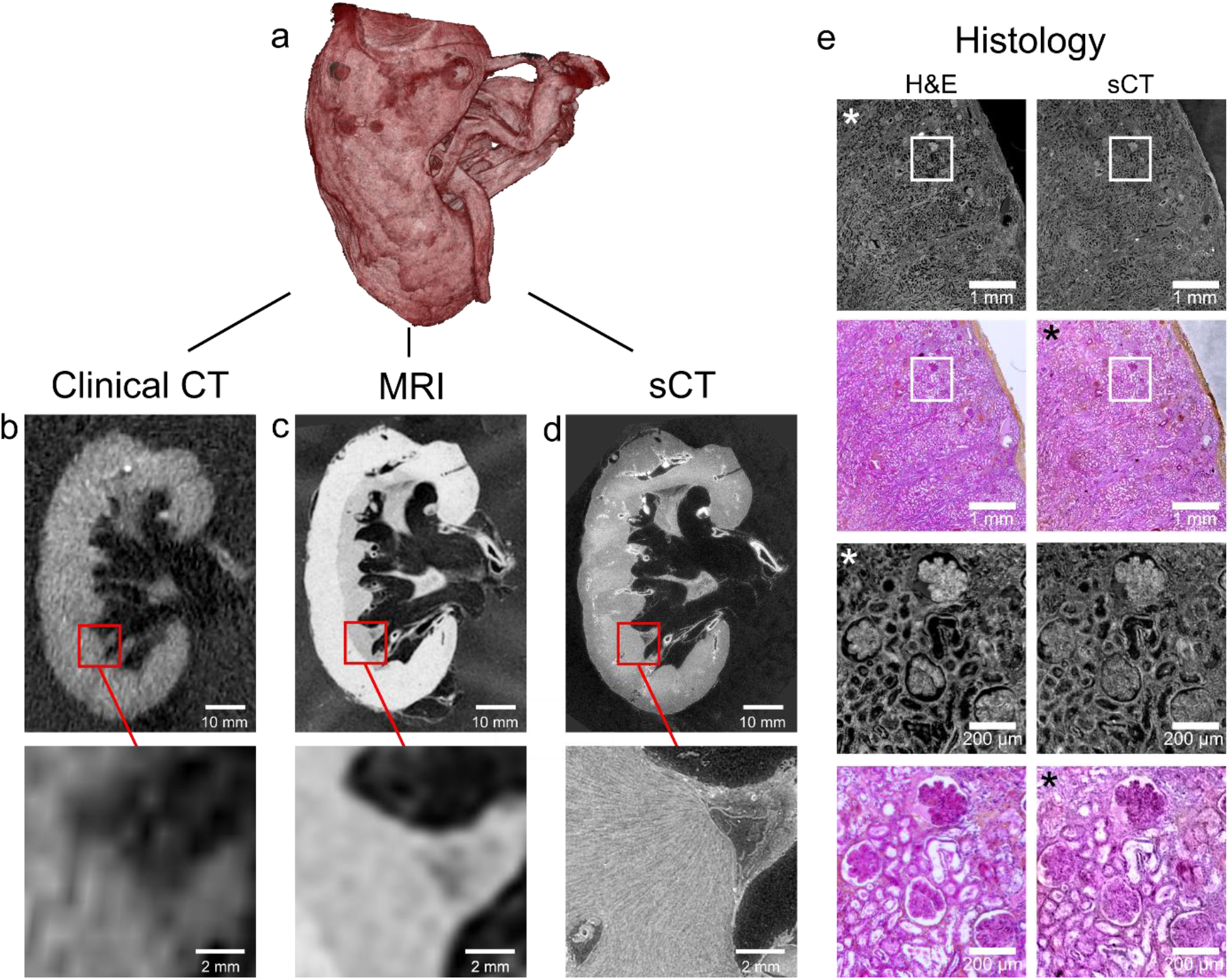
Visualization of an intact human kidney with clinical CT, MRI, sCT, and histology. **a**, 3D view of the human kidney. **b**, Cross-section of the kidney imaged with medical CT, with a voxel-size of 625 µm. **c**, Cross-section of the kidney imaged with a 3T medical MRI, with a voxel-size of 220×200×200 µm^3^. **d**, Cross-section of the kidney imaged with sCT, with a voxel-size of 25 µm. **e**, Comparison of sCT and a histopathological section stained with hematoxylin and eosin (H&E). Histology was performed after all other imaging modalities. The left column shows light micrography images of H&E-stained histopathological sections, the right column shows 2D sCT images. Images with a star on the top left have been pseudocolored (from sCT) or converted to gray levels before to be inverted in contrast (from histology). Modified from Walsh et al.^34^. All experiments followed the relevant governmental and institutional ethics regulations for human experiments.

Dehydration of the sample with ethanol increases the contrast for X-ray imaging, but significantly affects the MR images^52^. The majority of MR techniques are highly dependent on water content. Thus, by dehydrating the sample, the contrast between tissues that retain water well (e.g. adipose tissue) and those that dehydrate efficiently (e.g. muscle tissue) is enhanced. However, by decreasing the total water content, signal is reduced, and some techniques such as diffusion-weighted MR are no longer possible. One method of overcoming this problem is to not dehydrate the sample but to prepare the sample directly in a formalin mounting for performing MRI first. The contrast for sCT images is then reduced, but the signal-to-noise ratio of MRI images is increased. Although formalin- or paraformaldehyde-fixed tissue is the most used sample preparation method for ex-vivo MRI, this technique also affects the quality of the images^53^. The fixation of tissue with formalin or paraformaldehyde reduces T_1_, T_2_ and diffusivity of tissue, hence reducing the contrast-to-noise in images. One approach, is to wash the brain to remove fixative before MRI imaging which can restore the pre-fixation T_2_ values though not the T_1_^54^. Washing will also compromise the preceding degassing steps, so the relative importance of each imaging modality must be carefully weighed against the overall imaging pipeline and multi-modal registration requirements.

### Expertise needed to implement the protocol

The protocol outlined here requires prior experience in handling human or animal organs. If human organs are used, a trained clinician should be part of the study to handle body embalming and organ dissection. This protocol was developed and optimized for imaging large organs, a scientist who has experience in CT or sCT is required as imaging such large structure is not trivial and requires thorough preparation.

## Method

### Biological materials

- Human organs obtained from donated body **! CAUTION** Institutional and governmental ethics regulations concerning human experiment must be followed. All the experiments herein were authorized by the European Synchrotron Radiation Facility, the Laboratoire d’Anatomie des Alpes Françaises (LADAF) and the Hannover Institute of Pathology at Medizinische Hochschule, Hannover (ethics vote no. 9022_BO_K_2020). Transport and imaging protocols were approved by the Health Research Authority and Integrated Research Application System (HRA and IRAS) (200429) and the French Health Ministry. All organ dissections respected the memory of the deceased. The post-mortem study was conducted according to the Quality Appraisal for Cadaveric Studies scale recommendations^55^.

### Reagents

- 4% neutral buffered formalin, (50% Hygeco, cat. no. 07040 + 50% Hygeco, cat. no. 07020; diluted to 4%) **! CAUTION** Toxic and flammable. Handle with care, in a fume hood. Avoid inhalation and exposure to the skin or eyes.
- Ethanol 96% (Cheminol France, cat. no. 20×4-2)
- Agar-Agar (Nature et Aliments, cat. no. 510190) **! CRITICAL** The gelling properties of agar differ depending on the producer. We recommend using the supplier listed to reproduce the results presented here.
- Demineralized water (Sigma Aldrich, cat. no. 38796)

### Equipment

- Fume hood (Iberis laboratoire, cat. no. SPR18)
- Gloves Nitril (Showa, cat. no. 7500PF)
- Paper towels (Paredes, cat. no. 404857)
- Dissecting instruments (Fisher Scientific, cat. no. 11738551)
- Vacuum pump (Marshall scientific, cat. no. VME8)
- Vacuum dessicator (SP Bel-Art, cat. no. F42400-4031)
- Fridge (Fisher Scientific, cat. no. 22651264)
- Large leak-proof container in PET (Medline Scientific, cat. no. 129-0592, or Lock & Lock, cat. no. INL-403, or Lock & Lock, cat. no. INL-203)
- Magnetic vortex system (Fisherbrand, cat. no. 15349654)
- Electric grater (Seb, cat. no. DO201141)
- Ladle (Tefal, cat. no. K1180214)
- Syringe 50ml (PentaFerte, cat. no. 002022960)
- Sieve (Profilstore, cat. no. toilinox_PGRI1ME213)
- Tape (3M, cat. no. 764)
- Drill (DeWALT, cat. no. DCD791P2-QW)
- Synchrotron microtomography setup. We used the BM05 beamline of the ESRF. See Walsh et al^34^ for detailed information on the sCT setup.

### Procedure

#### Organ procurement

**!CAUTION** Any experiments involving the use of human organs must be ethically approved by the relevant institutional or governmental committees. Experiments were performed in accordance with a protocol approved by the the European Synchrotron Radiation Facility, the Laboratoire d’Anatomie des Alpes Françaises (LADAF) and the Hannover Institute of Pathology at Medizinische Hochschule, Hannover (ethics vote no. 9022_BO_K_2020). Transport and imaging protocols were approved by the Health Research Authority and Integrated Research Application System (HRA and IRAS) (200429) and the French Health Ministry.

1. Collect the organ of interest either by extracting it from a donated body (option A) or through a biobank, a surgery, or a transplantation (option B).
  A. **Embalming of the body and organ extraction – Timing ∼3 d**
    i. (Optional) Embalm the body after death by injecting sequentially 4,500ml formalin diluted to 1.15% in a solution containing lanolin and 4,500ml formalin diluted to 1.44% into the right carotid artery (performed by a licenced medical practitioner). **!CAUTION** Formalin is volatile and highly toxic. It is an eye, respiratory and skin irritant and a probable carcinogen. Work inside a chemical fume hood in a well-ventilated area and wear appropriate personal protective equipment (gloves, eye protection, and laboratory coat). **PAUSE POINT** The body can be stored up to 2 years if fixed.
    ii. Extract the organs of interest from the body.
    iii. Remove surrounding fat and connective tissue.
  B. **Organ procurement from a biobank – Timing depending on the administrative and delivery delays**
    i. Retrieve the organ from a biobank, a surgery, or a transplantation. **Organ fixation – Timing 4 d**
2. (Optional) Fix completely the organ by immersing it in 4% neutral buffered formalin at room temperature. We used 4 days minimum for fixation by formalin to comply with safety rules linked to autopsies in case of pathogenic agent (such as covid-19). **Preparation of the organ before mounting** **!CAUTION** Degassing with a vacuum pump should be carried out under a fume hood or in a room equipped with a ventilation system of sufficient capacity to avoid inhalation of formalin or ethanol gases.
3. (Optional) Wash the sample with tap water to remove the fixative agent.
4. Prepare the organ for mounting by dehydration using multiple baths of ethanol and degassing it under vacuum (option A). With the vacuum degassing protocol, damage could be seen in some fragile organs (typically the brain). Thus, an alternative protocol to prepare the sample without vacuum degassing was developed using thermal cycles (option B). The ethanol can be replaced with 4% formalin solution to avoid the dehydration of the sample where this is undesirable e.g. MR imaging. Modifications to this procedure may be necessary as timing and number of cycles depends heavily on the composition and size of the organ. A better optimization of this step could be achieved by further investigation. ? TROUBLESHOOTING
  A. **Dehydration and vacuum degassing – Timing 16 d**
    i. Immerse the organ in a bath of pre-degassed 50% ethanol. The solution must be at least 4x the volume of the organ.
    ii. Place the container with the organ inside a desiccator and remove the free and dissolved gas in the tissue by diminishing the pressure using a diaphragm vacuum pump. **!CRITICAL STEP** The degassing should be performed in cycles of increase duration. Each cycle is typically 15min to 30min for a human organ, such as a heart or a kidney. For each cycle, the vacuum pumping is stopped after bubbling decreases or if it becomes too strong in intensity and could potentially damage the organ. Each time a new cycle starts, the time before first bubble apparition should increase. The degassing time has to be adapted to each organ depending on its composition and size. There is no typical time, it has to be chosen empirically by looking at the bubbling regime. This will strongly depend on the vacuum pump and desiccator quality. ? TROUBLESHOOTING
    iii. Wait until equilibrium (Our tests show that typically 4 days for a human organ such as lung, heart, or brain are sufficient for each ethanol concentration. With 2 days only, we saw incomplete equilibrium visible as density gradients in the scans).
    iv. Repeat steps (i), (ii), and (iii) with three successive baths of pre-degassed ethanol at 60%, 70%, and finally 70% with a degassing between each bath.
    v. Perform a final degassing of the organ for few minutes just before mounting it with the agar agar crushed gel in the mounting jar.
  B. **Dehydration and thermal cycles degassing (alternative protocol for fragile organs) – Timing 5 weeks**
    i. Performed at least 4 thermal cycles by immersing the organ in four successive baths of highly degassed 50%, 60%, 70%, 70%, ethanol. Close the container without entrapping bubbles, then store at 4°C during 5 days, and bring back to room temperature for each cycle. The solution has to be at least 4x the volume of the organ. **PAUSE POINT** Once at 70% of ethanol, the organ can be stored at room temperature for months to years before continuing with the mounting steps of the protocol. **Preparation of the organ mounting gel - Timing 2d** The following produces 5 L of agar gel:
5. Boil 5L of demineralised water in a container with a magnetic vortex system.
6. Once above 80 degrees, slowly pour 100g of agar agar (20g/L) powder in the water and keep the vortex until good dissolution.
7. Once dissolved, stop the agitation and pour the liquid in a suitable container.
8. Once gelation has been achieved (∼12h depending on the temperature), cut the agar gel in cubes of approximately 2×2×2cm^3^.
9. Immerse the cubes in 13,75 L of 96% ethanol. The volume ratio is chosen to ensure a final ethanol concentration of 70%.
10. Wait until the density of the blocks is close to the one of ethanol (∼24h).
11. Check the equilibrium of the solution by agitating the solution to put the gel cubes in suspension and ensure that they sink slowly to the bottom of the container (over several dozen of seconds).
12. Degas the solution using a vacuum pump and a desiccator for 2 cycles of approximately one hour to avoid cracking the agar cubes.
13. Store a third of the degassed cubes, in an air-tight container.
14. Crush the remaining cubes using an electric grater.
15. Store in an airtight container for future use, and degas again just before use. **Organ mounting and degassing – Timing ∼3h**
16. Fill the bottom of the leak-proof cylindrical container with agarose cubes a few centimeters.
17. Fill half the container with crushed gel. **!CRITICAL STEP** Use a ladle with careful slow movements when manipulating the agar solution to avoid increasing the dissolved gas in the solution or entrapment of bubbles.
18. Carefully immerse the organ in the gel and set it to the desired position.
19. Cover the organ with crushed agar.
20. Degas the whole container to remove entrapped bubbles. **!CRITICAL STEP** This degassing steps only aims at removing entrapped bubbles. As all the components were degassed before this should limit the amount of dissolved gas. This vacuum degassing should be done only with short pumping times (2-3 minutes) for several cycles to help remove the visible bubbles. **!CRITICAL STEP** If necessary, gentle tapping on the desiccator can help bubbles travel to the top
21. Add more of the agar-ethanol solution.
22. Use a sieve to press on the agar gel from the top to compact it around the organ. Use a syringe to remove the excess ethanol from on top of the sieve, then add more agar and ethanol. **!CRITICAL STEP** Be careful as once the agar is compact, the bubbles cannot rise to the top, and therefore degassing can no longer be carried out on this part of the container. All movements have to be done slowly and carefully to ensure not to entrap bubbles when compacting the agar.
23. If necessary, Push the agar down in the container with your fingers to compact the agar around the sample and maintain the sample in position until it cannot move anymore in the jar. **!CRITICAL STEP** A space must be left between the sample and the container wall to avoid border effects during X-ray imaging.
24. Degas the whole container using a vacuum pump to remove the gas added during the last 3 steps and to avoid trapping bubble in the agar gel.
25. Repeat the last 4 steps until the agar is compact enough below, around and above the sample to avoid any movement of the specimen. An example of insufficiently compacted agar is shown in the online supplementary video 1. ? TROUBLESHOOTING
26. Fill the leak-proof container to the top with the agar-ethanol solution.
27. Drill a small hole of few millimeters of diameter in the center of the lid.
28. Screw the lid on the container. If the container is not directly leak-tight, apply latex on the container thread to ensure a good seal.
29. After sealing, check that all air was removed from the container by applying a slight pressure on the lid and ensuring that ethanol comes out of the hole. **!CRITICAL STEP** Agar can block the hole and prevent ethanol from coming out, in which case use a needle to clear the hole
30. Apply a piece of tape onto the hole on the lid to avoid gas exchange between the inside and outside of the container, and to act as a safety exhaust in case of bubbling event during scanning.
31. Apply tape around the lid to protect it and prevent it from being accidentally opened.
32. Assess that no bubbles remain in the container. Once the agar is compact, the bubbles cannot travel easily to the top. If, on inspection, few small bubbles are trapped, wait for 24h. As all the components were degassed before mounting, most of the bubbles will naturally dissolve. If not sufficient and few small bubbles are still present, an overnight refrigeration may resorb them. If this does not work, the mounting must be restarted from the beginning. **!PAUSE POINT** The mounted organ can be stored at room temperature for months to years before imaging, and can be imaged many time without any intervention as long as there is no bubbling event due to too high X-ray dose or significant movement of the sample. **3D imaging of the mounted sample (Variable depending on the imaging technique)**
33. Image the mounted sample with sCT, µCT, clinical CT, or MRI. The sample can be imaged with as many of these imaging techniques as desired. ? TROUBLESHOOTING
34. (Optional) Perform histological analysis on the biological sample

## ? Troubleshooting

| Step | Problem | Possible reasons | Possible solutions |
| --- | --- | --- | --- |
| 4 | After the dehydration, a significant shrinkage of the organ is observed | The tissue composition makes it particularly sensitive to ethanol dehydration | A gentler transition is required. Increase the number of ethanol baths to smooth the transition of the organ to 70% ethanol, or use lower final concentration |
| 4A(ii) | During degassing, the organ starts to float on the ethanol, thus dehydrating and damaging a part of the sample. | Air is trapped inside the organ and the decrease in pressure makes the bubbles grow | Stop the pump as soon as this phenomenon occurs. Gently squeeze out the trapped air and restart the vacuum degassing. The sample can also be forced into the ethanol by pressing from above with a grid to ensure that it cannot reach the surface of the liquid |
| 25 | After the mounting, the sample embedded in the agar gel still moves inside the container when it is rotated (see Supplementary Video 1 | The agar gel is not compact enough to hold the sample well, and/or there are not enough agar cubes at the bottom of the jar. | If there are agar cubes at the bottom of the jar, repeat steps 22-25 until the sample no longer moves under gentle agitation. If there are no cubes, then dismount, |
|  | online). This lack of rigidity can create artefacts in the images and prevent registration of scans if multiple ROI are imaged |  | put the cubes and restart the mounting procedure. An alternative solution would be to use jars with internal protruding structure at their bottom that would block rotation of the crushed agar |
| 33 | Formation of bubbles in the container during the scan reducing the quality of the images | The sample was not degassed enough and the amount of dissolved gas remaining in the solution was too high<br>The dose applied to the sample was too high | Check the scanning parameters before making any new scan. Preserve the sample in the fridge for one night to try to resorb the bubbles. If this solution is ineffective, try a rapid vacuum degassing without dismounting after having opened the jar, put an extender and put ethanol above the agar gel to allow efficient escape of the bubbles and filling of the cavities when putting back the atmospheric pressure. If not sufficient, the mounting has to be redone |

### Timing

Step 1, Organ procurement: ∼3 d

Step 2, Fixation of the organ: 4 d

Step 3-4A or 3-4B, preparation of the organ before mounting: 16–25 d

Step 5-15, preparation of the organ mounting gel: 2d

Step 16-32, organ mounting and degassing: ∼3h

Step 33-34, Imaging and histology: Variable depending on the imaging technique and setup

### Anticipated results

This protocol provides a method to prepare, and stabilize large organs or biological samples for imaging at high resolution using sCT, µCT, clinical CT, or MRI imaging. Typical images from human organs are shown in Figure 2 and Figure 5. This sample preparation procedure provides high contrast images (depending on the imaging modality), prevents sample movement during scanning, is compatible with multiple imaging modalities, and preserves the morphological characteristics of the tissue compared to other sample preparation methods like paraffin-embedding. The sample can be stored at least one year without movement or deterioration. This method enables the 3D investigation of large biological structures like human organs without damaging them. These high-resolution images can provide qualitative and quantitative information on the healthy or pathological behavior of an organ, for instance the effect of the COVID-19 in human lungs^34^.

## Supporting information

SV1: Example of insufficient compact agar

## Data Availability

Image data used to create the figures present in this protocol paper are publicly available from the ESRF data repository (https://human-organ-atlas.esrf.eu) or from the corresponding authors.

## Supplementary information

Supplementary Video 1

Example of insufficient compact agar (MP4 7836 kb)

## Acknowledgements

We thank S. Bayat (INSERM) for help during the test phase, P. Masson (LADAF) for dissections of donors’ bodies, H. Reichert (ESRF) and R. Tori for general support of the project and C. Muzelle, R. Homs, C. Jarnias, F. Cianciosi, P. Vieux, P. Cook, L. Capasso and A. Mirone for their help in setup developments and improvements. We also thank R. Engelhardt, A. Muller Brechlin, C. Petzold, N. Kroenke, and M. Kuhel for help with histology and autopsies. This project has been made possible in part by grants number 2020-225394 from the Chan Zuckerberg Initiative DAF, an advised fund of Silicon Valley Community Foundation, The ESRF - funding proposals md1252 and md1290, the Royal Academy of Engineering (CiET1819/10) and the MRC (MR/R025673/1). M.A. acknowledges grants from the National Institutes of Health (HL94567 and HL134229) This work was supported by the German Registry of COVID-19 Autopsies (DeRegCOVID, www.DeRegCOVID.ukaachen.de; supported by the Federal Ministry of Health - ZMVI1-2520COR201), and the Federal Ministry of Education and Research as part of the Network of University Medicine (DEFEAT PANDEMIcs, 01KX2021).

## Author contributions

P.T., P.D.L., D.D.J., M.A., C.L.W. and W.L.W. conceptualized the project and designed experiments. M.A., C.W., P.T., A.B., C.L.W., and J.B. performed autopsies and sample preparation. P.T. designed and built instrumentation and performed HiP-CT imaging; S.M. designed sample holders; P.T. designed and implemented tomographic reconstruction methods; J.B., P.T., P.D.L. wrote the paper. All authors assisted in reviewing and revising the manuscript.

